## Supplemental materials for "Molecular portraits of colorectal cancer morphological regions"

### Supplemental material for “Molecular portraits of morphological regions of colorectal cancer”

#### Supplemental methods

##### Samples

This retrospective cross-sectional study used tumor samples from patients with CRC who were examined at Masaryk Memorial Cancer Institute, Brno, Czech Republic in years 2002-2015. The study was approved by the Committee for Ethics of Masaryk Memorial Cancer Institute, Brno, Czech Republic (number 2018/861/MOU). Written informed consent was obtained from all study participants in accordance with the Declaration of Helsinki. Inclusion criteria for this study were: age > 18 years, clinical and histopathologically confirmed diagnosis of primary CRC. Standard clinical and histopathological variables (TNM, grade etc.) were retrieved for all patients. Failure of laboratory analyses (problematic sample preparation, low quality and/or quantity of isolated RNA, low quality of expression data) was a reason for excluding these samples from the study.

##### Gene expression profiling

The RNA extraction was performed from formalin-fixed paraffin-embedded histopathological slides using AllPrep® DNA/RNA Kits (Qiagen, Hilden, Germany) according to their specific manufacturer's instructions. A few modifications were made to the protocol: FFPE slides (2x 3µm) were bathed in a solution to remove paraffin (3x in xylene for 5 min and 3x in ethanol for 5 min). Tumor tissue was spotted with 8ul PKD puffer and collected from slides using a scalpel. Purification was done for total RNA, including small RNAs. For elution, 20ul RNA free water (1 min. incubation) was used and then repeated with eluate. The extracted RNA served as input for a GeneChip™ WT Pico Reagent Kit (Thermo Fisher Scientific, Waltham, MA, USA) for analysis of the transcriptome on whole-transcriptome arrays. We selected the input amount from the recommended range according to the manufacturer’s instructions. Total RNA from HeLa cells provided in the kit was used as a positive control together with a high-quality low-concentration RNA isolated from a serum as a low input control. Clariom™ D Array for human samples (Thermo Fisher Scientific, Waltham, MA, USA) was used for target hybridization to capture both coding and multiple forms of non-coding RNA. Finally, the arrays were scanned using Affymetrix GeneChip™ Scanner 3000 7G (Thermo Fisher Scientific, Waltham, MA, USA). The sample preparation and analysis were performed according to the manufacturer ́s instructions. The protocol included several control points in which the workflow was monitored. All the samples complied with the quality control requirements and none of the samples were excluded from the analysis.

##### Data preprocessing

All resulting CEL files were processed using `Bioconductor` (1)(v.3.15) packages `oligo` (2)(v.1.60), `affycoretools` (v1.68) and, for Clariom D chip annotation, `pd.clariom.d.human` (v.3.14). For the quality control we used `AffyPLM` (v.147) and imposed a maximal median Normalized Unscaled Standard Errors (NUSE) of 1.12. In all, n=202 passed all the quality control steps and were normalized together using `RMA` (`oligo`) with core-probeset summarization. Further, the array data was summarized at gene level by selecting the most variable probeset per unique `EntrezID` and entries corresponding to missing `HUGO` symbols, speculative transcripts, and short non-coding RNA were discarded resulting in a reduced list of 27,302 unique genes. All data analyses were performed in R 4.2 (3). Batch effects were removed using `ComBat` (4) from package `sva` (v.3.44.0)

#### Supplemental Figure 1

Morphotype distribution per case (unique tumor) and intersections thereof: some cases had several morphotypes profiled.

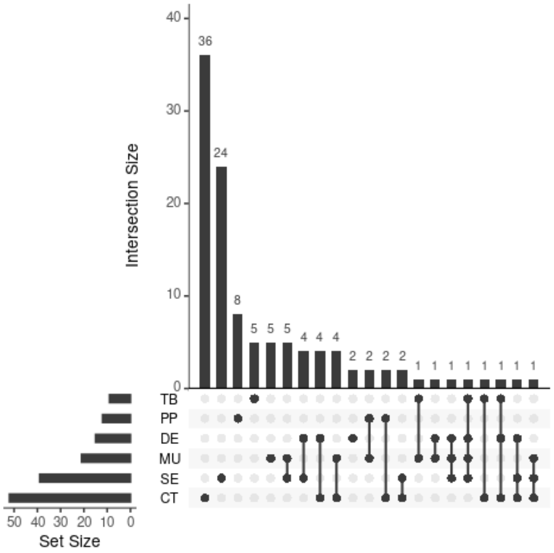

#### Supplemental Figure 2

SF2 A: Epithelial signatures from Pelka et al., 2021. Only statistically significant scores (NES) are shown.

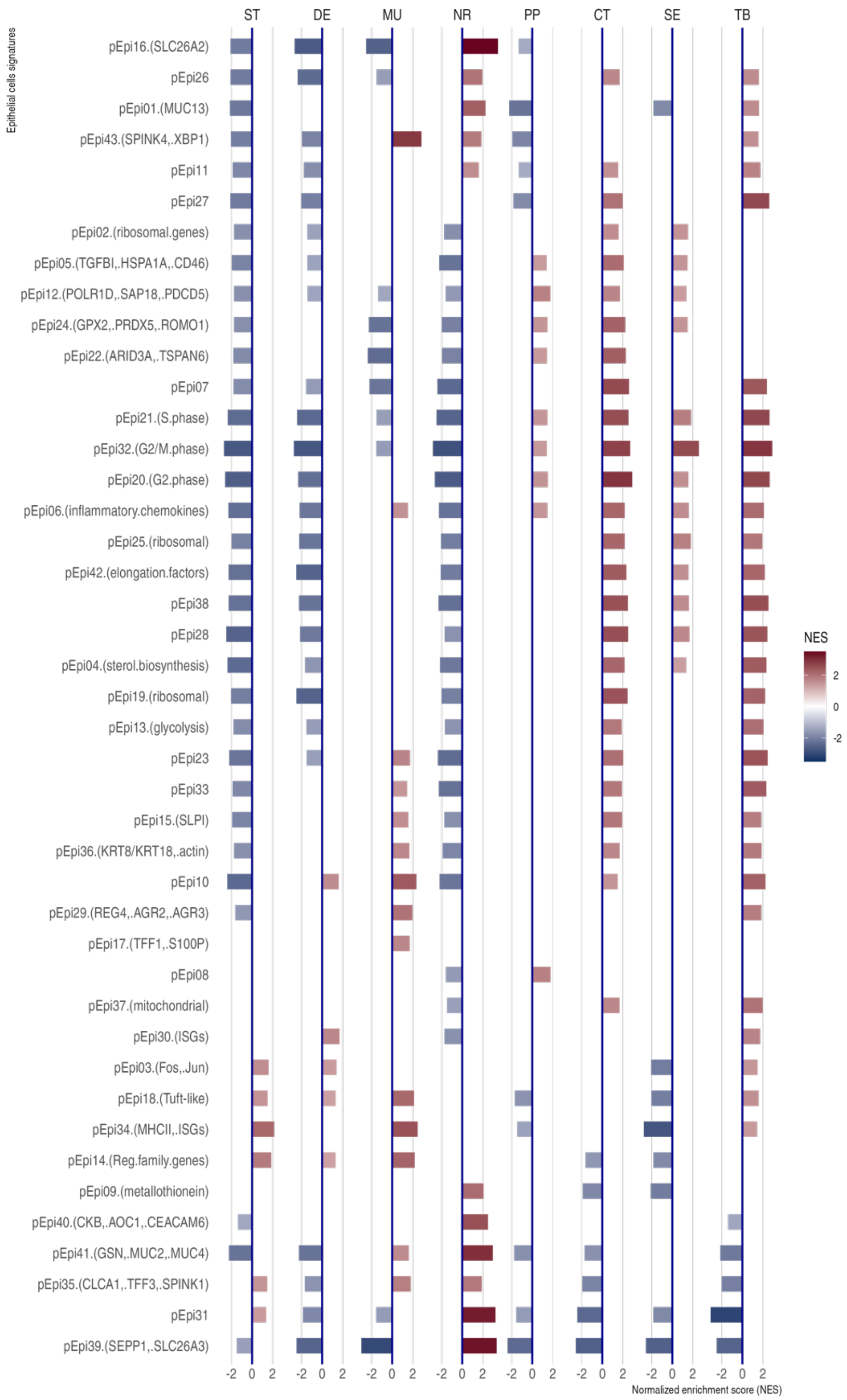

SF2 B: immune signatures from Pelka et al., 2021. Only statistically significant scores (NES) are shown.

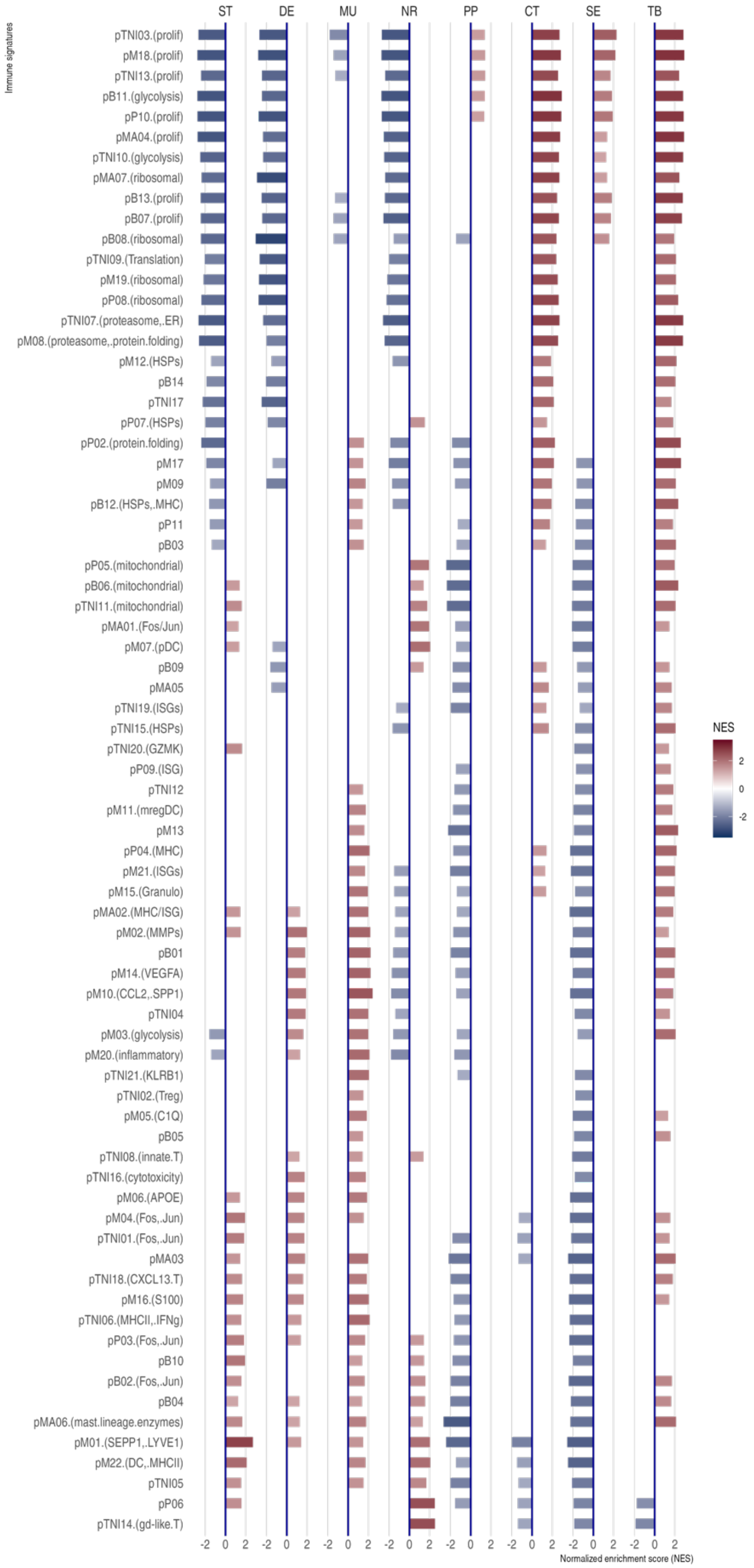

SF2 C: Stromal signatures from Pelka et al., 2021. Only statistically significant scores (NES) are shown.

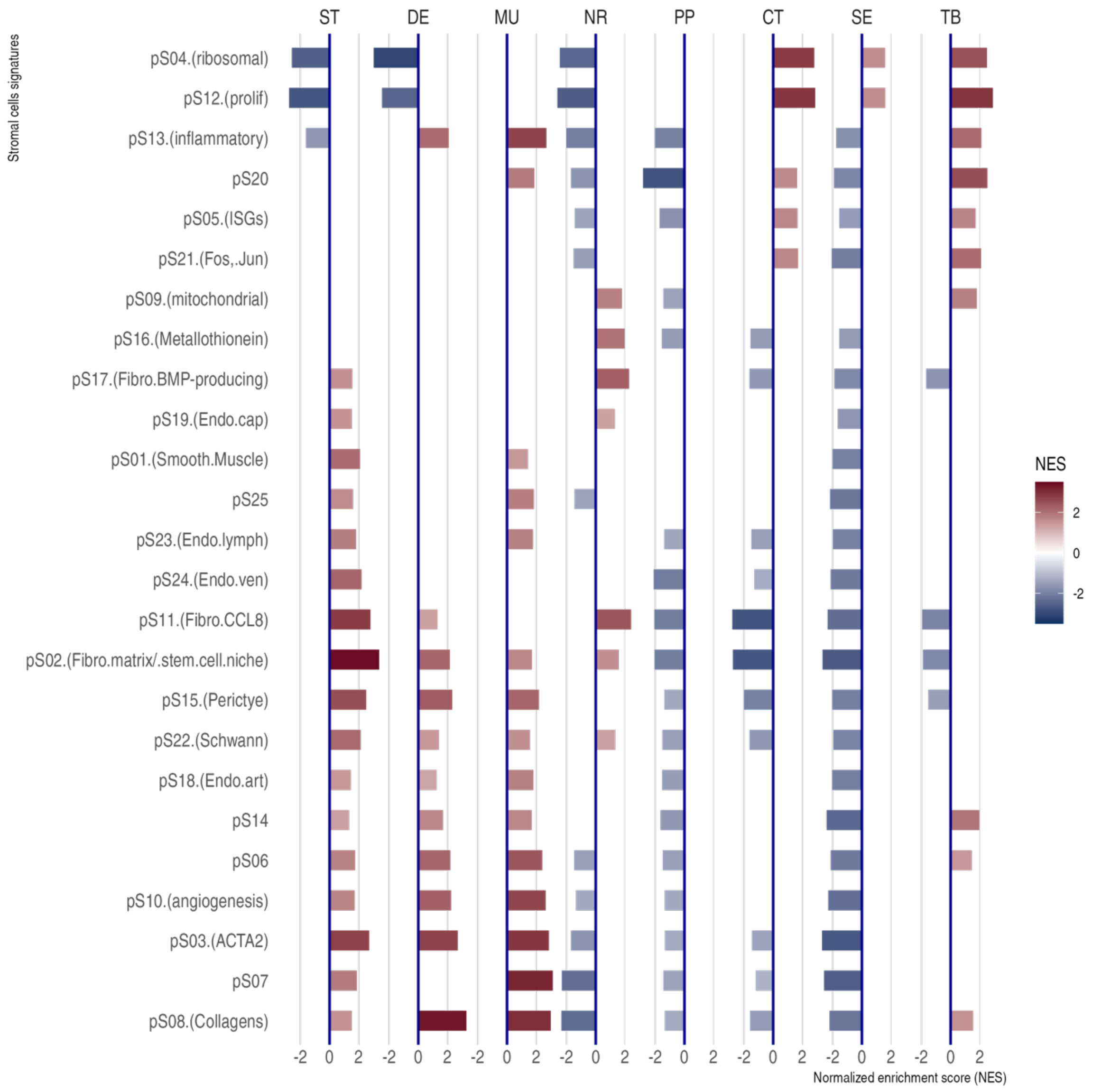

#### Supplemental Figure 3

Principal component analysis of hallmark pathways GSEA scores: loadings for the first two principal components, i.e., contribution of pathways to the first two axes.

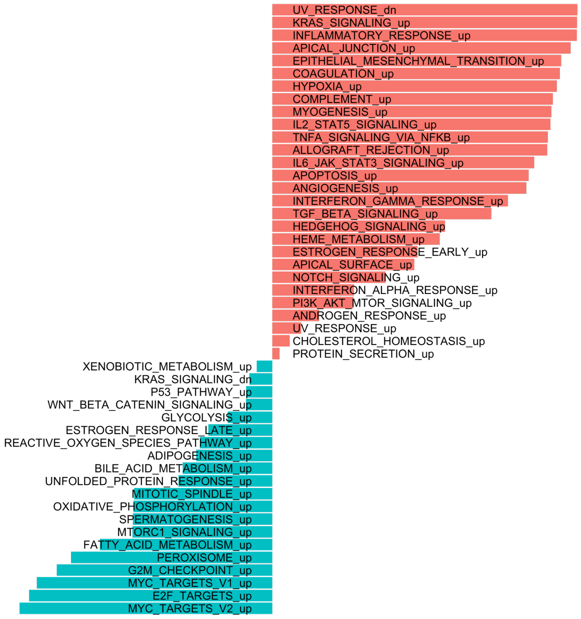


1st principal component loadings (40.6% of total variation)

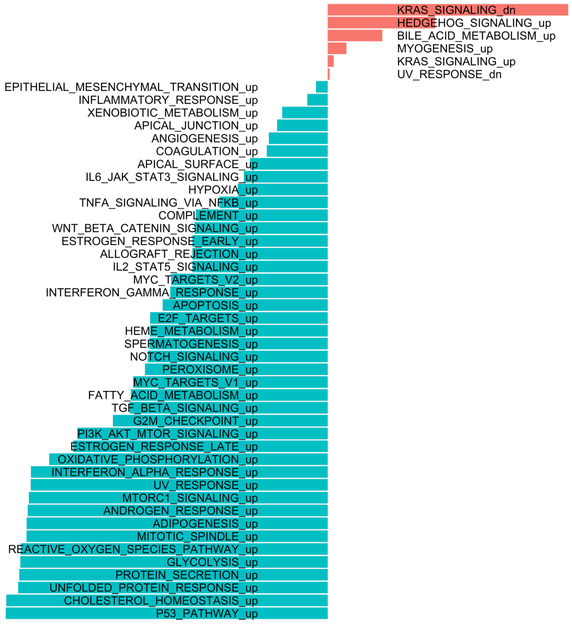


2nd principal component loadings (33.5% of total variation)

#### Supplemental Figure 4

Hallmark pathways differential activation between pairs of morphotypes. Here we compare the results from GSEA applied to differentially expressed genes between pairs of morphotypes originating from all cases to results of GSEA applied to differentially expressed genes between pairs of morphotypes originating from the same section (tumor) (i.e., matched pairs of morphotypes). All results are shown, including the statistically not significant one. First four columns correspond to pairs of morphotypes from all cases, while the last four to matched pairs of morphotypes.

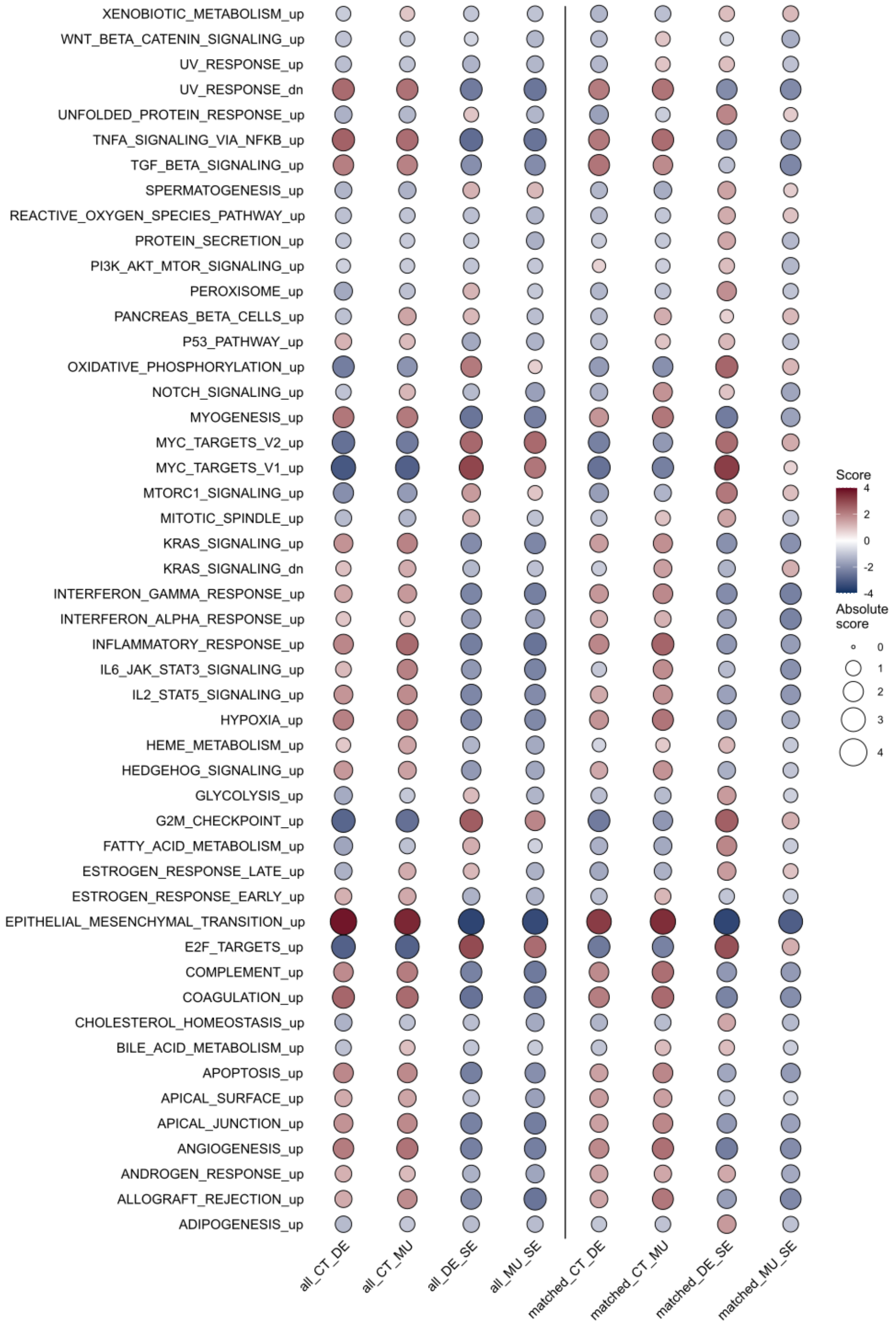

### Supplemental Figure 5

SF5 A

**Case P1949**

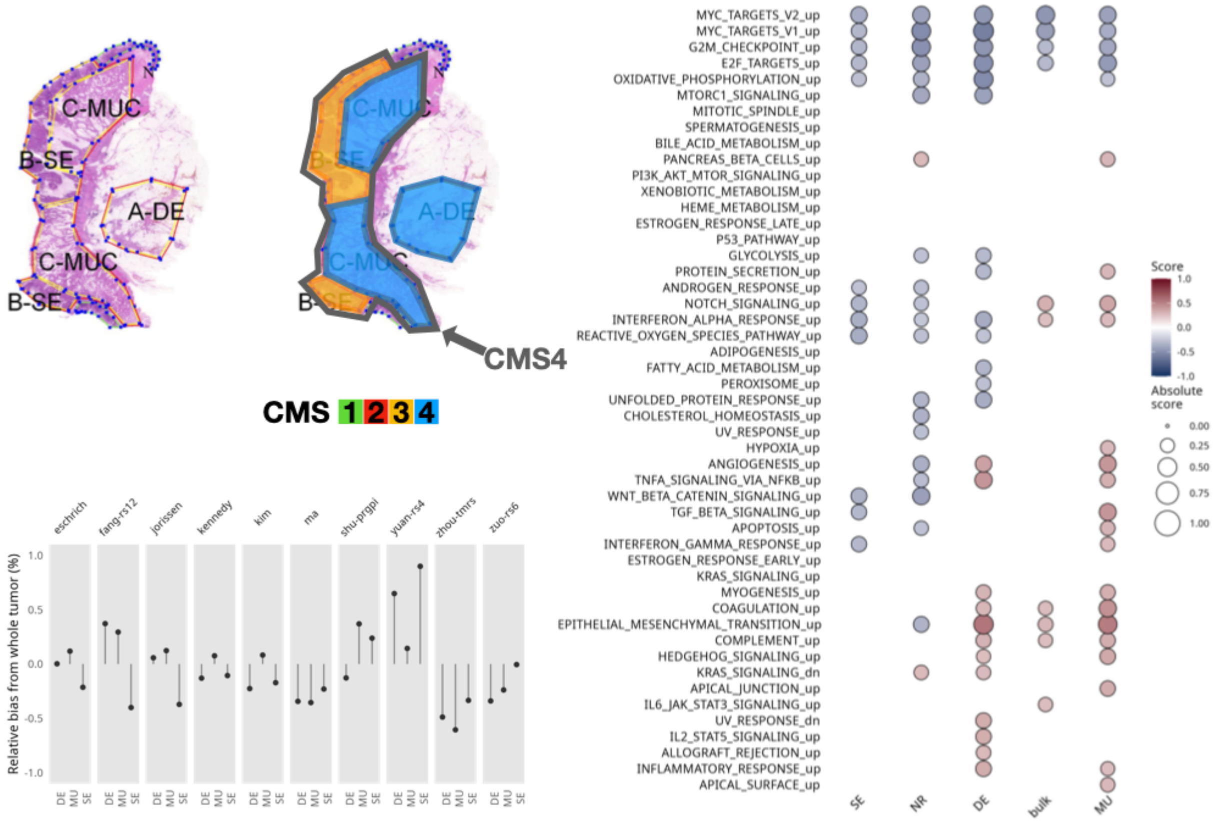


Survival scores

Hallmark pathways (GSVA scores)

SF5 B

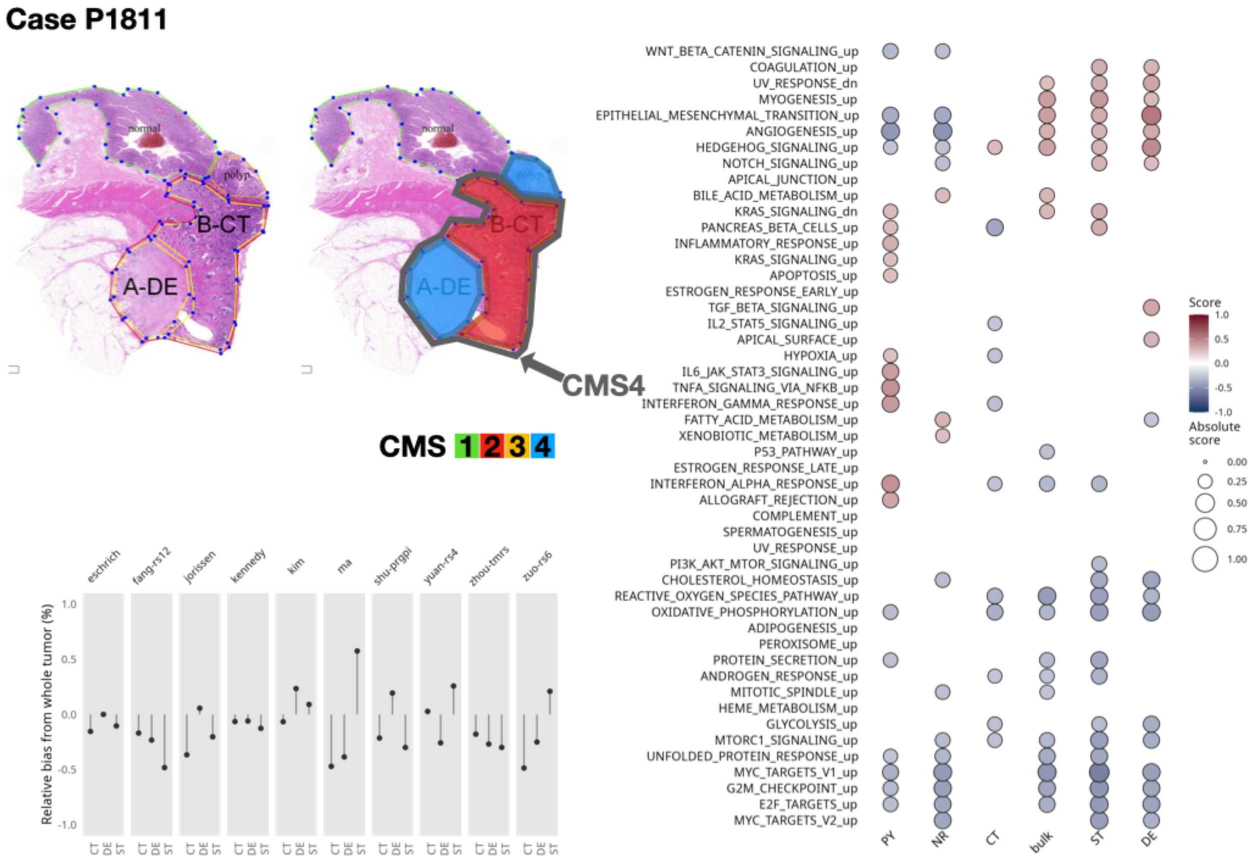


Survival scores

Hallmark pathways (GSVA scores)

#### Supplemental Figure 6

SF6 A: Normalized enrichment scores from GSEA for selected resistance signatures (from C2 section of MSigDB). Only significant scores are shown.

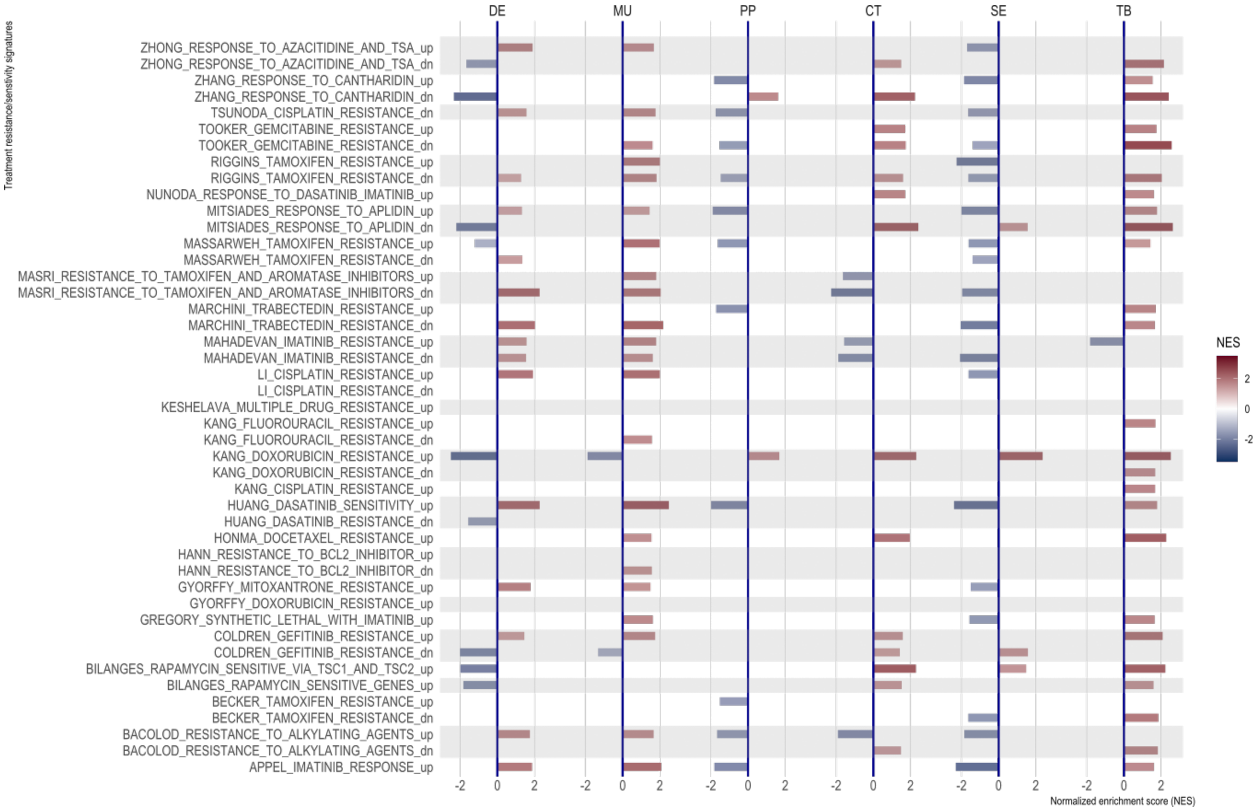

SF6 B: Resistance scores (GSVA) per patient and morphotype for a number of cases where the whole-tumor prediction is contradicted by some regional score.

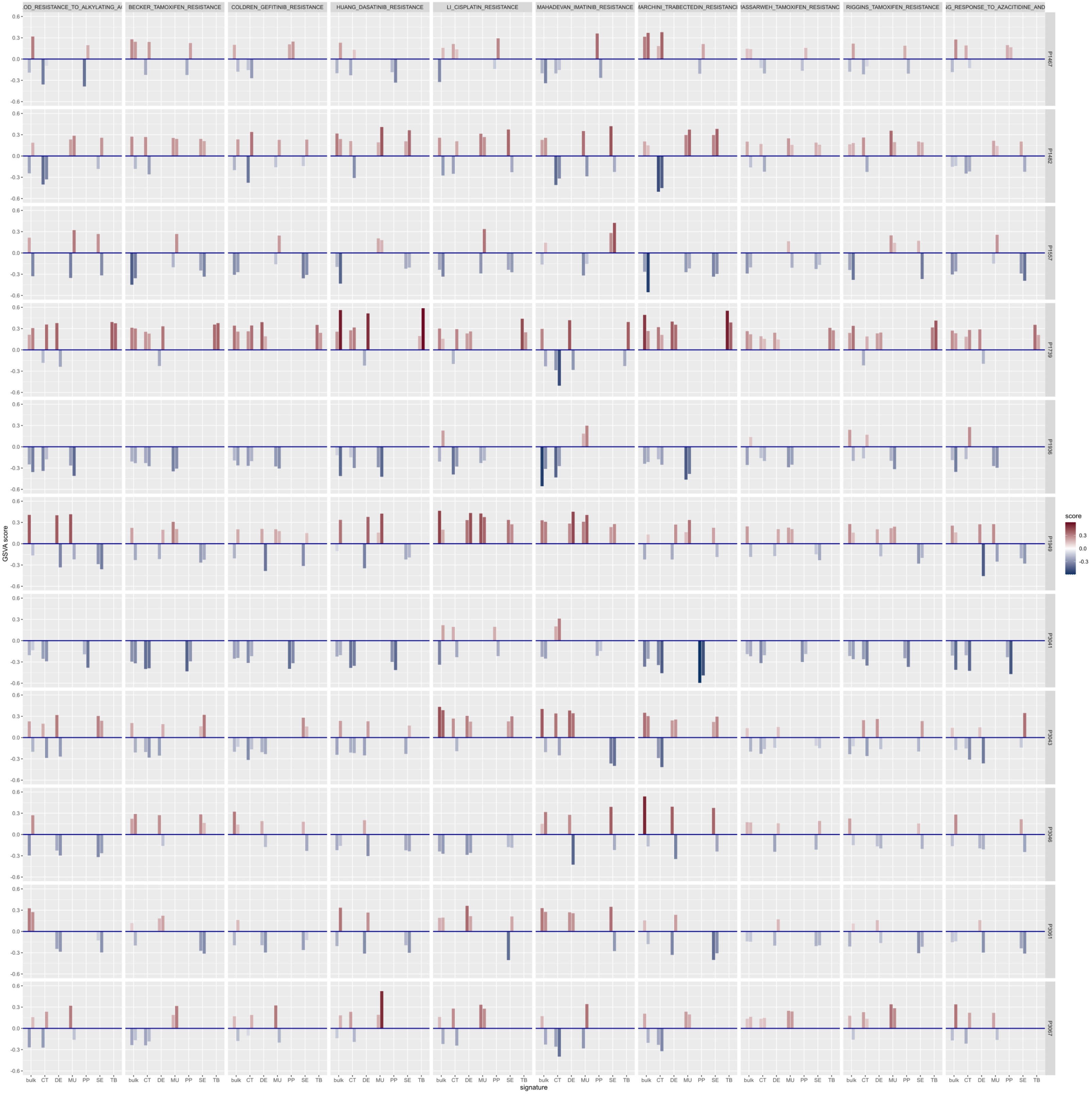

#### Supplemental Figure 7

Molecular subtypes and morphotypes in all samples, including non-core samples. Note that the sets of samples in A and C are the same as in Figure 4, as only core samples had also at least two distinct morphological regions.

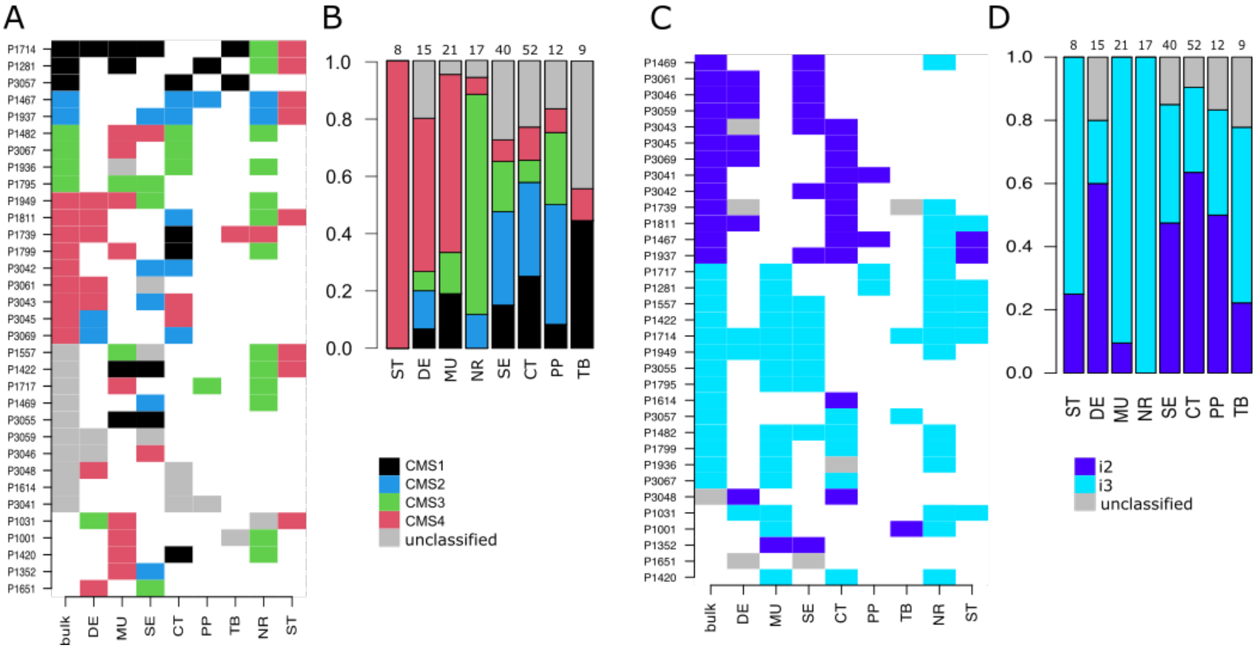

#### Supplemental Table 1

Main clinical parameters of the study cohort.

| | Overall (N=111) |
|---|---|
| **gender** | |
| F | 58 (52.3%) |
| M | 53 (47.7%) |
| **pT** | |
| pT1 | 1 (0.9%) |
| pT2 | 6 (5.4%) |
| pT3 | 96 (86.5%) |
| pT4 | 8 (7.2%) |
| **pN** | |
| pN0 | 61 (55.0%) |
| pN1 | 32 (28.8%) |
| pN2 | 18 (16.2%) |
| **M** | |
| M0 | 91 (82.0%) |
| M1 | 20 (18.0%) |
| **grade** | |
| 1 | 11 (9.9%) |
| 2 | 52 (46.8%) |
| 3 | 36 (32.4%) |
| Missing | 12 (10.8%) |
| **stage.cat** | |
| II | 59 (53.2%) |
| III | 32 (28.8%) |
| IV | 20 (18.0%) |
| **site.cat** | |
| left | 32 (28.8%) |
| right | 54 (48.6%) |
| transversum | 23 (20.7%) |
| Missing | 2 (1.8%) |

#### Supplemental Table 2

Distribution of main clinical parameters per morphotype (and tumor-adjacent normal and supportive stroma).

| | CT (N=52) | DE (N=15) | MU (N=21) | PP (N=12) | SE (N=40) | TB (N=9) | NR (N=17) | ST (N=8) | Overall (N=174) |
|---|---|---|---|---|---|---|---|---|---|
| **Sex** | | | | | | | | | |
| F | 26 (50.0%) | 8 (53.3%) | 12 (57.1%) | 6 (50.0%) | 20 (50.0%) | 8 (88.9%) | 11 (64.7%) | 6 (75.0%) | 97 (55.7%) |
| M | 26 (50.0%) | 7 (46.7%) | 9 (42.9%) | 6 (50.0%) | 20 (50.0%) | 1 (11.1%) | 6 (35.3%) | 2 (25.0%) | 77 (44.3%) |
| **pT stage** | | | | | | | | | |
| pT1 | 1 (1.9%) | 0 (0%) | 0 (0%) | 0 (0%) | 0 (0%) | 0 (0%) | 0 (0%) | 0 (0%) | 1 (0.6%) |
| pT2 | 2 (3.8%) | 1 (6.7%) | 0 (0%) | 1 (8.3%) | 2 (5.0%) | 1 (11.1%) | 0 (0%) | 0 (0%) | 7 (4.0%) |
| pT3 | 45 (86.5%) | 13 (86.7%) | 18 (85.7%) | 11 (91.7%) | 37 (92.5%) | 7 (77.8%) | 16 (94.1%) | 8 (100%) | 155 (89.1%) |
| pT4 | 4 (7.7%) | 1 (6.7%) | 3 (14.3%) | 0 (0%) | 1 (2.5%) | 1 (11.1%) | 1 (5.9%) | 0 (0%) | 11 (6.3%) |
| **pN stage** | | | | | | | | | |
| pN0 | 30 (57.7%) | 10 (66.7%) | 12 (57.1%) | 6 (50.0%) | 25 (62.5%) | 4 (44.4%) | 10 (58.8%) | 5 (62.5%) | 102 (58.6%) |
| pN1 | 18 (34.6%) | 2 (13.3%) | 6 (28.6%) | 3 (25.0%) | 9 (22.5%) | 2 (22.2%) | 5 (29.4%) | 2 (25.0%) | 47 (27.0%) |
| pN2 | 4 (7.7%) | 3 (20.0%) | 3 (14.3%) | 3 (25.0%) | 6 (15.0%) | 3 (33.3%) | 2 (11.8%) | 1 (12.5%) | 25 (14.4%) |
| **M stage** | | | | | | | | | |
| M0 | 43 (82.7%) | 13 (86.7%) | 18 (85.7%) | 10 (83.3%) | 33 (82.5%) | 7 (77.8%) | 13 (76.5%) | 6 (75.0%) | 143 (82.2%) |
| M1 | 9 (17.3%) | 2 (13.3%) | 3 (14.3%) | 2 (16.7%) | 7 (17.5%) | 2 (22.2%) | 4 (23.5%) | 2 (25.0%) | 31 (17.8%) |
| **Grade** | | | | | | | | | |
| 1 | 1 (1.9%) | 1 (6.7%) | 3 (14.3%) | 1 (8.3%) | 9 (22.5%) | 0 (0%) | 2 (11.8%) | 1 (12.5%) | 18 (10.3%) |
| 2 | 25 (48.1%) | 4 (26.7%) | 6 (28.6%) | 6 (50.0%) | 19 (47.5%) | 2 (22.2%) | 5 (29.4%) | 2 (25.0%) | 69 (39.7%) |
| 3 | 18 (34.6%) | 3 (20.0%) | 10 (47.6%) | 4 (33.3%) | 6 (15.0%) | 6 (66.7%) | 10 (58.8%) | 5 (62.5%) | 62 (35.6%) |
| Missing | 8 (15.4%) | 7 (46.7%) | 2 (9.5%) | 1 (8.3%) | 6 (15.0%) | 1 (11.1%) | 0 (0%) | 0 (0%) | 25 (14.4%) |
| **Stage** | | | | | | | | | |
| II | 29 (55.8%) | 10 (66.7%) | 12 (57.1%) | 6 (50.0%) | 23 (57.5%) | 4 (44.4%) | 9 (52.9%) | 4 (50.0%) | 97 (55.7%) |
| III | 14 (26.9%) | 3 (20.0%) | 6 (28.6%) | 4 (33.3%) | 10 (25.0%) | 3 (33.3%) | 4 (23.5%) | 2 (25.0%) | 46 (26.4%) |
| IV | 9 (17.3%) | 2 (13.3%) | 3 (14.3%) | 2 (16.7%) | 7 (17.5%) | 2 (22.2%) | 4 (23.5%) | 2 (25.0%) | 31 (17.8%) |
| **Tumor site** | | | | | | | | | |
| left | 18 (34.6%) | 3 (20.0%) | 5 (23.8%) | 1 (8.3%) | 14 (35.0%) | 1 (11.1%) | 3 (17.6%) | 1 (12.5%) | 46 (26.4%) |
| right | 21 (40.4%) | 9 (60.0%) | 14 (66.7%) | 8 (66.7%) | 17 (42.5%) | 6 (66.7%) | 12 (70.6%) | 6 (75.0%) | 93 (53.4%) |
| transversum | 12 (23.1%) | 2 (13.3%) | 2 (9.5%) | 2 (16.7%) | 8 (20.0%) | 2 (22.2%) | 2 (11.8%) | 1 (12.5%) | 31 (17.8%) |
| Missing | 1 (1.9%) | 1 (6.7%) | 0 (0%) | 1 (8.3%) | 1 (2.5%) | 0 (0%) | 0 (0%) | 0 (0%) | 4 (2.3%) |

#### Supplemental Table 3

Table of gene expression signatures: see ST3-other_sigs.xlsx

#### Supplemental Table 4

Table of GSEA scores (NES) for “other” signatures: see ST4-GSEA_res_all_signif_other.xlsx (see also ST3 for signatures).

#### Supplemental Table 5

List of differentially expressed genes (limma tables) per morphotype in contrast with pooled profile. See ST5-limma_toptable_per_morphotype.xlsx

#### Supplemental Table 6

GSEA results for genes in ST5, for whole MSigDB collection: ST6-GSEA_res_full-MSigDB.xlsx

#### Supplemental Table 7

List of differentially expressed genes (limma tables) per morphotype in contrast with all other five morphotypes: ST7-limma_toptable_morphotype_vs_rest.xlsx

#### Supplemental Table 8

GSEA results for genes in ST7, for whole MSigDB collection: ST8-GSEA_res_all_signif_other-morphotype_vs_rest-6morphos.xlsx

#### Supplemental Table 9

List of differentially expressed genes (limma tables) per matched pairs of morphotypes: ST9-limma_toptable_matched_pairs.xlsx

#### Supplemental Table 10

GSEA results for genes in ST9, for whole MSigDB collection: ST10-GSEA_res_all_signif_matched_pairs.xlsx

#### Supplemental Table 11

List of differentially expressed genes (limma tables) for pairs of morphotypes: ST11-limma_toptable_morphotype_pairs.xlsx

#### Supplemental Table 12

GSEA results for genes in ST11, for whole MSigDB collection: ST12-GSEA_res_morphotype_pairs_signif.xlsx

#### Supplemental Table 13

List of prognostic signatures tested.

| Gene signature ID | Reference |
|---|---|
| **Eschrich** | Steven Eschrich, Ivana Yang, Greg Bloom, et al. *Molecular Staging for Survival Prediction of Colorectal Cancer Patients.* Journal of Clinical Oncology 2005 23:15, 3526-3535 |
| **Jorissen** | Jorissen RN, Gibbs P, Christie M, et al. *Metastasis-Associated Gene Expression Changes Predict Poor Outcomes in Patients with Dukes Stage B and C Colorectal Cancer.* Clin Cancer Res. 2009 |

| | |
|---|---|
| | Dec 15;15(24):7642-7651. doi: 10.1158/1078-0432.CCR-09-1431. PMID: 19996206; PMCID: PMC2920750. |
| **Kennedy** | Kennedy RD, Bylesjo M, Kerr P, et al. *Development and independent validation of a prognostic assay for stage II colon cancer using formalin-fixed paraffin-embedded tissue*. J Clin Oncol. 2011 Dec 10;29(35):4620-6. doi: 10.1200/JCO.2011.35.4498. Epub 2011 Nov 7. PMID: 22067406. |
| **Kim** | Kim, SK., Kim, SY., Kim, C.W. *et al. A prognostic index based on an eleven gene signature to predict systemic recurrences in colorectal cancer*. *Exp Mol Med* **51**, 1–12 (2019). https://doi.org/10.1038/s12276-019-0319-y |
| **Ma** | Ma Xiao-Bo, Xu Yuan-Yuan, Zhu Meng-Xuan, Wang Lu. *Prognostic Signatures Based on Thirteen Immune-Related Genes in Colorectal Cancer.* Frontiers in Oncology, 10, 2021, doi: 10.3389/fonc.2020.591739 |
| **Yuan-rs4** | Yuan, Yihang et al. *Development and Clinical Validation of a Novel 4-Gene Prognostic Signature Predicting Survival in Colorectal Cancer*. Frontiers in Oncology. 10, 2020, doi: 10.3389/fonc.2020.00595 |
| **Zhou-tmrs** | Zhou R, et al. *A robust panel based on tumour microenvironment genes for prognostic prediction and tailoring therapies in stage I-III colon cancer*. EBioMedicine. 2019 Apr;42:420-430. doi: 10.1016/j.ebiom.2019.03.043. Epub 2019 Mar 24. PMID: 30917936; PMCID: PMC6491960. |
| **Shu-prgpi** | Shu P, Wu J, Tong Y, Xu C, Zhang X. Gene pair based prognostic signature for colorectal colon cancer. Medicine (Baltimore). 2018 Oct;97(42):e12788. doi: 10.1097/MD.0000000000012788. PMID: 30334969; PMCID: PMC6211904. |
| **Fang-rs12** | Fang Z, Xu S, Xie Y, Yan W. *Identification of a prognostic gene signature of colon cancer using integrated bioinformatics analysis.* World J Surg Oncol. 2021 Jan 13;19(1):13. doi: 10.1186/s12957-020-02116-y. PMID: 33441161; PMCID: PMC7807455. |
| **Zuo-rs6** | Zuo S, Dai G, Ren X. Identification of a 6-gene signature predicting prognosis for colorectal cancer. Cancer Cell Int. 2019 Jan 5;19:6. doi: 10.1186/s12935-018-0724-7. PMID: 30627052; PMCID: PMC6321660. |
